# State-dependent activity dynamics of hypothalamic stress effector neurons

**DOI:** 10.1101/2022.02.10.479856

**Authors:** Aoi Ichiyama, Samuel Mestern, Gabriel B. Benigno, Kaela E. Scott, Brian L. Allman, Lyle Muller, Wataru Inoue

## Abstract

The stress response necessitates an immediate boost in vital physiological functions from their homeostatic operation to elevated emergency response. However, neural mechanisms underlying this state-dependent change remain largely unknown. Using a combination of *in vivo* and *ex vivo* electrophysiology with computational modeling, we report that corticotropin releasing hormone (CRH) neurons in the paraventricular nucleus of the hypothalamus (PVN), the effector neurons of hormonal stress response, rapidly transition between distinct activity states through recurrent inhibition. Specifically, *in vivo* optrode recording shows that under non-stress conditions, CRH_PVN_ neurons often fire with rhythmic brief bursts (RB), which, somewhat counterintuitively, constrains firing rate due to long (∼2 s) inter-burst intervals. Stressful stimuli rapidly switch RB to continuous single spiking (SS), permitting a large increase in firing rate. A spiking network model shows that recurrent inhibition can control this activity-state switch, and more broadly the gain of spiking responses to excitatory inputs. In biological CRH_PVN_ neurons *ex vivo*, the injection of whole-cell currents derived from our computational model recreates the *in vivo*-like switch between RB and SS, providing a direct evidence that physiologically relevant network inputs enable state-dependent computation in single neurons. Together, we present a novel mechanism for state-dependent activity dynamics in CRH_PVN_ neurons.

## INTRODUCTION

Mounting an optimal stress response is critical for survival. This requires neural mechanisms that effectively switches vital physiological functions between homeostatic operation and emergency response. For example, the activation of the hypothalamic-pituitary-adrenal (HPA) axis elevates the systemic levels of glucocorticoids, a hallmark of stress. Under non-stress conditions, glucocorticoids are also produced, albeit at lower levels, for their widespread physiological actions (de Kloet et al., 2005; Lightman, 2008). While HPA axis (patho)physiology has been extensively studied with relevance to stress-related disorders (Gold and Chrousos, 1999; McEwen, 2005), we know surprisingly little about the basic neural mechanisms that powerfully and bidirectionally transition HPA axis activity from baseline to levels of elevated stress.

The brain ultimately drives the HPA axis by exciting a group of neuroendocrine neurons that synthesize corticotropin releasing hormone (CRH). The cell bodies of CRH neurons reside in the paraventricular nucleus of the hypothalamus (PVN) and project their axons to the median eminence. At the median eminence, axonally released CRH enters the hypophyseal portal circulation and initiates the downstream hormonal cascades (Swanson and Sawchenko, 1980). Numerous studies have shown that various acute stressors induce *de novo* expression of immediate early genes in CRH_PVN_ neurons, establishing their stress-induced activity increase in the time scale of minutes to hours (Cole and Sawchenko, 2002; Ek et al., 2000; Itoi et al., 2004; Matovic et al., 2020; Wamsteeker Cusulin et al., 2013). Recent *in vivo* fibre photometry imaging revealed highly dynamic activity changes where population Ca^2+^ signals in CRH_PVN_ neurons (a proxy for population activity) increase rapidly (within seconds) and reversibly in response to repetitive, short aversive stimuli (Daviu et al., 2020; Kim et al., 2019a, 2019b; Yuan et al., 2019).

How do individual CRH_PVN_ neurons fire action potentials during the rapid transitions between baseline and stress conditions? Although the *in vivo* firing activity of single-units in the PVN was first studied more than 30 years ago (Hamamura et al., 1986; Kannan et al., 1987; Saphier and Feldman, 1985, 1988, 1991), these pioneering experiments lacked neurochemical identification; *i.e.*, it could not be confirmed whether the spiking activity was of CRH neurons or another sub-population of neurons within the PVN. This is because *in vivo* extracellular electrophysiology is inherently blinded for neuronal identities, and CRH neurons are intermingled with a mosaic of neuron-types deep in the hypothalamus (Simmons and Swanson, 2009). Thus, despite decades of research on CRH_PVN_ neurons, their firing activities *in vivo* have remained inaccessible. This is an important gap in our knowledge because single-neuron firing patterns result from dynamic interaction between intrinsic properties and network dynamics (Destexhe et al., 2001; Herz et al., 2006; Leng and MacGregor, 2018), and hence will provide essential clues to probe how CRH_PVN_ neurons−the gatekeepers of stress outputs−process the converging brain commands during baseline conditions and in response to stress.

Here we have implemented *in vivo* single-unit extracellular recording of optogenetically-identified CRH_PVN_ neurons in mice, and subsequently developed a network model that explains *in vivo* state-dependent firing dynamics changes. For the first time, we report that CRH_PVN_ neurons fire in two distinct modes: rhythmic brief bursts (RB) and single spikes (SS) of random spike intervals. Somewhat counterintuitively, RB occurred under the baseline conditions and constrained the overall firing rate at low levels due to long (∼2 s) inter-burst silence. On the other hand, sustained SS underlies stress-induced increase in firing. Using tight combinations of experimental and computational approaches, we show that a recurrent inhibitory network generates RB and constrains the overall firing rate due to negative feedback inhibition. A spiking network model showed that release from recurrent inhibition upon stress stimuli permits continuous SS that increases the overall firing rate, thereby revealing a novel mechanism that enables efficient bidirectional controls of stress output neurons. To validate the spiking network model, we used *ex vivo* patch-clamp electrophysiology and injected excitatory and inhibitory inputs received by a cell in the computational model into actual biological neurons. CRH_PVN_ neurons in these experiments exhibited *in vivo*-like bursting behavior when driven by the network current, but not when driven by simple step-current stimuli. Ultimately, our collective results identified a novel network mechanism that underlies state-dependent firing activity dynamics at the effector neurons of hormonal stress response.

## RESULTS

### Identifying CRH neurons *in vivo*

To examine firing patterns of individual CRH_PVN_ neurons *in vivo*, we used optrode recording (Lima et al., 2009). Briefly, we first “tagged” CRH neurons by expressing an excitatory opsin channelrhodopsin2 (ChR2) using Cre-dependent adeno-associated virus (AAV) injected into the PVN of CRH-Ires-Cre mice crossed with Ai14 td-tomato reporter line (Wamsteeker Cusulin et al., 2013). This resulted in ChR2-EYFP expression in CRH_PVN_ neurons that express td-tomato (**Fig. 1A**), similar to recent studies using the same approach (Bittar et al., 2019; Füzesi et al., 2016; Kim et al., 2019a; Wamsteeker Cusulin et al., 2013; Yuan et al., 2019). Important for the desired PVN neuron identification, these ChR2-expressing neurons could now be detected on the basis of their light-induced spike firing. We isolated 36 single-units in the PVN area from 10 adult male mice under urethane anaesthesia. Among these pre-isolated single units, 18 neurons (from 10 mice) were “light-responsive”; *i.e.*, they fired action potentials in response to pulses of blue light (5 ms, λ = 465 nm) with short latency (7.2 ± 2.6 ms, n = 18; **Fig. 1B and C**. This responsiveness to light had a binary effect (**Fig. 1D**): the light-responsive neurons reliably increased their probability of firing (7.1 ± 6.3 % pre-onset to 66.3 ± 21.3 % post-onset, n = 18), whereas the light-non-responsive neurons did not (3.3 ± 3.1 % pre-onset to 2.0 ± 2.8 % post-onset, n = 18). Furthermore, in response to longer light pulses (50 ms), the light-responsive neurons fired a train of action potential with frequency adaptation (**Fig. 1E, F**). Consequently, these light-responsive neurons were defined as CRH_PVN_ neurons.

**Figure 1.**
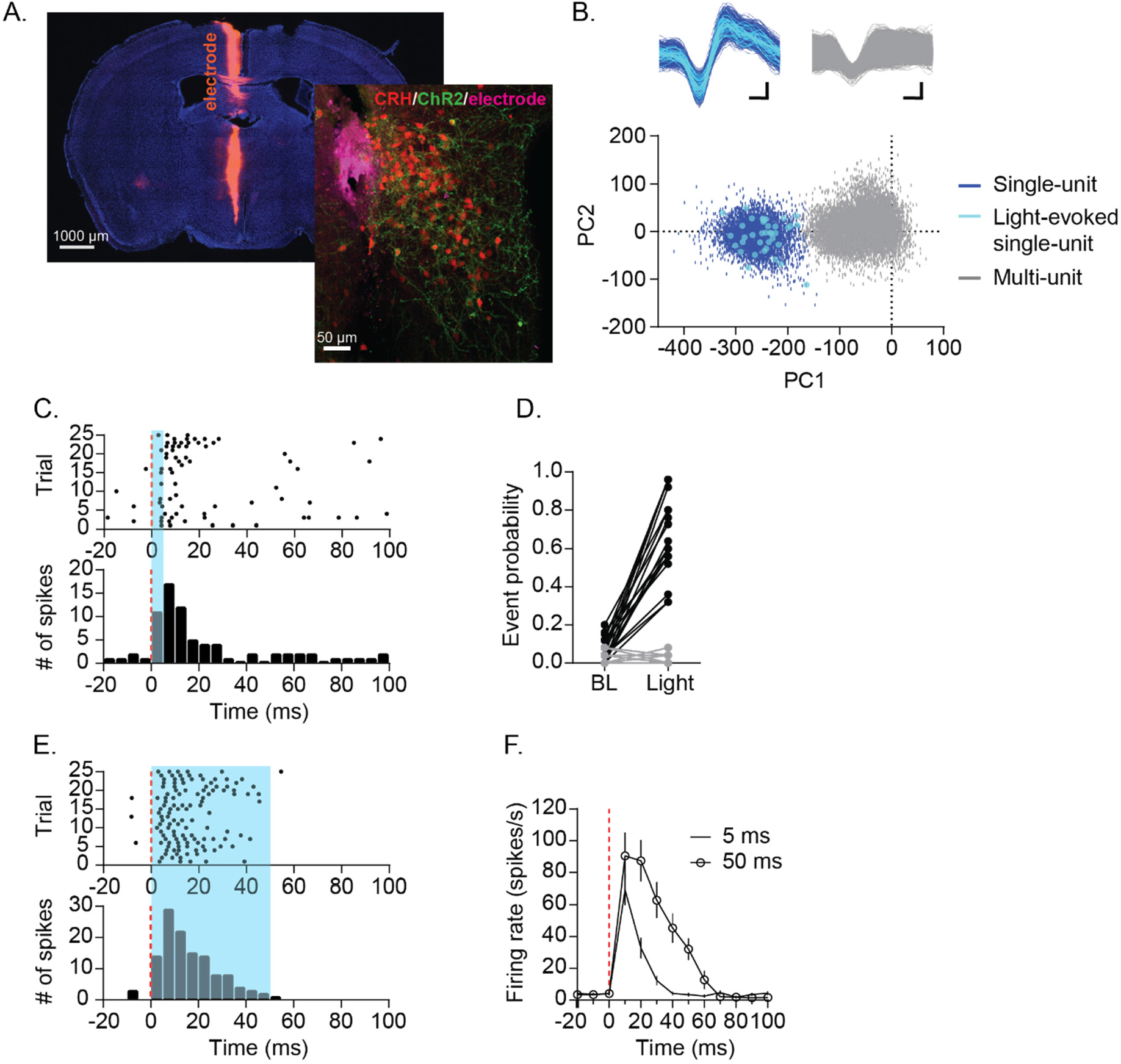
Optogenetic identification of CRH_PVN_ neuron single-unit. **A.** Electrode tract (purple), TdTomato-expressing CRH_PVN_ neurons (red) and ChR2-EYFP (green) expression. **B.** An example for an isolated single-unit (blue) that also responded to light (cyan). **C.** Raster plot (top) and peristimulus time histogram (PSTH, bottom) in response to 5 ms blue light from a representative unit. **D.** A summary graph for the probability of firing before (-20 to 0 ms) and after blue light illumination (0 to 20 ms). Light-responsive units in black (n = 18) and non-responsive units in grey (n = 18). **E.** Raster plot (top) and PSTH (bottom) for a representative single-unit responding to 50 ms blue light. **F.** Summary of firing rate time course following 5 ms (tick) and 50 ms (open circle) blue light. n = 18. Standard deviation is represented as error bars.

### CRH_PVN_ neurons can fire in two distinct modes

We found that, under the baseline (non-stress) conditions, many CRH_PVN_ neurons occasionally fired in a distinct brief train of high-frequency (>100 Hz) bursts, in addition to spikes with variable inter-spike-intervals (ISI, **Fig 2A**). **Figure 2B** shows an ISI histogram of a representative neuron that demonstrated a bimodal distribution with a sharp peak around 2-10 ms (burst) and another wider peak around 100 ms (single spikes, SS). Burst trains typically showed the shortest ISI at the start and followed by a few spikes with slightly increased ISIs. Thus, to quantify these brief burst firing, we set a criterion for burst detection with ISI ≤6 ms for the start of a burst, and subsequent spikes were considered part of the burst train as long as ISI remained below 20 ms. Using this burst-detection criterion, we found that the majority of CRH_PVN_ neurons fired a brief burst at least once during the baseline (16 out of 18), but the burst rate was variable among CRH_PVN_ neurons (**Fig. 2C**). Figure 2-Figure Supplement 1 shows ISI histograms for all units. For the analysis of burst properties below, we used neurons that showed at least 60 bursts during 10 min baseline recording (> 0.1 bursts/s, purple circles in **Fig. 2C**, n = 10). Among these bursting CRH_PVN_ neurons, each individual burst episode was brief, on average 3.1 ± 0.5 spikes, ranging between 2–6 spikes (**Fig. 2D**). These brief bursts occurred at slow rhythms (rhythmic brief burst, RB), intervened by long, mostly silent, inter-burst intervals (IBI, 1.8 ± 0.6 s, n = 10, **Fig. 2E**).

**Figure 2.**
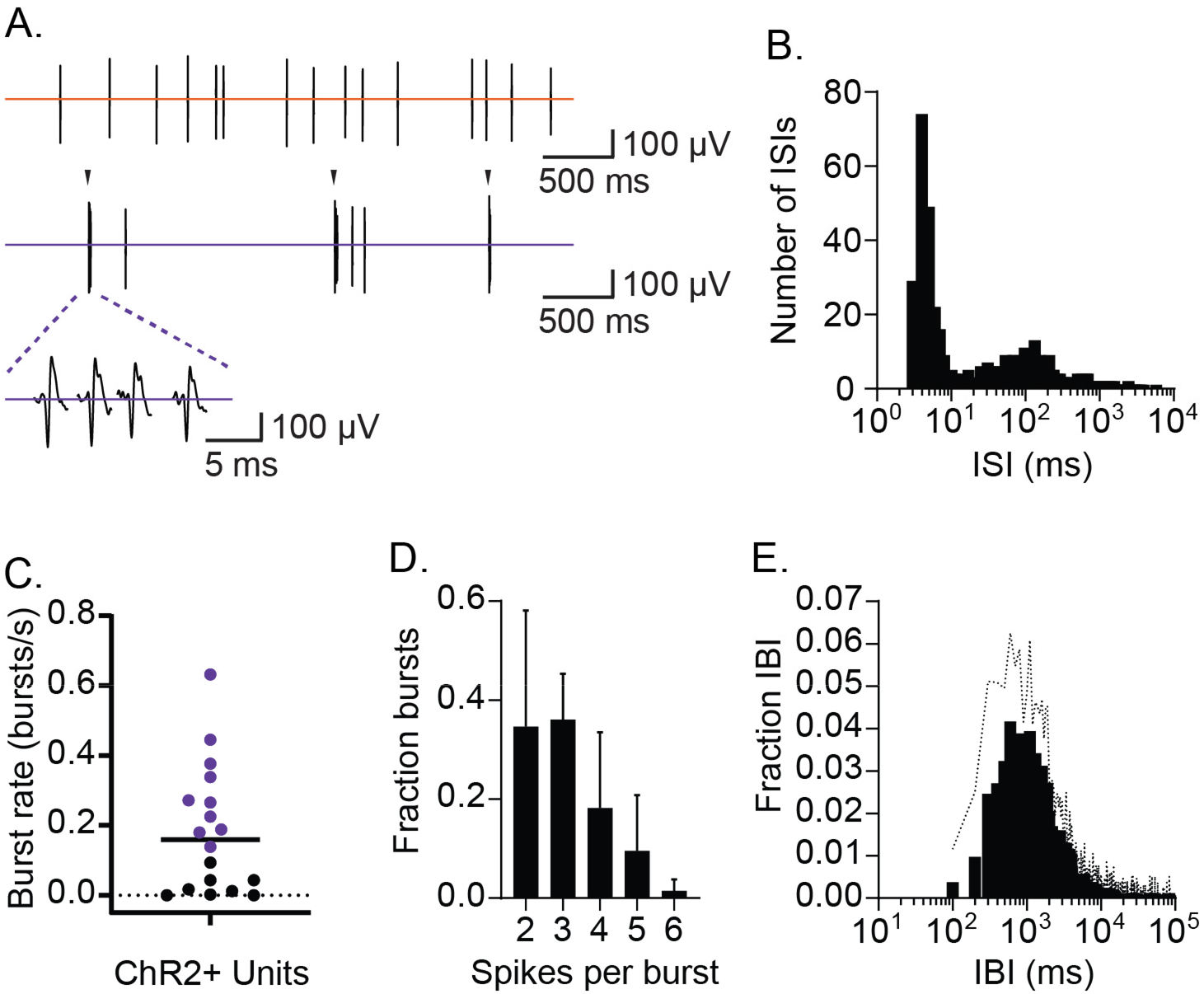
CRH_PVN_ neurons fire in rhythmic bursts and single spikes. **A.** Two distinct spike patterns in a representative single-unit (top: single spikes; bottom: bursts indicated by arrow). **B.** Inter-spike interval (ISI) distribution for the representative single-unit shown in A. **C.** A summary plot for burst rates among CRH_PVN_ neurons during non-stress baseline recordings. The criterion for burst firing neurons (purple) was set as 0.1 burst/s. A horizontal bar indicates the average. **D.** A summary graph for burst length distribution (n = 10). **E.** Summary inter-burst interval (IBI) distribution (n = 10). Standard deviation is represented in the graphs as the error bars (D) and dotted line (E).

### CRH_PVN_ neurons are constrained to low activity during rhythmic bursting

In many neurons, burst firing is postulated to carry behaviourally-relevant information, representing specific network states different from that represented by single spikes of variable frequencies (Krahe and Gabbiani, 2004; Steriade et al., 1993). The classic role of CRH_PVN_ neurons is to respond to stress stimuli with an increase in firing activity, driving the neuroendocrine release of CRH (Ulrich-Lai and Herman, 2009). Thus, we examined potential roles of RB in CRH_PVN_ neurons’ response to stress. To this end, we used electric stimulation of the sciatic nerve to model noxious sensory stimuli, as this approach has been shown to effectively elicit time-locked firing responses in median eminence-projecting parvocellular PVN neurons (Day et al., 1985; Hamamura et al., 1986; Kannan et al., 1987; Saphier, 1989; Saphier and Feldman, 1985, 1991) as well as to increase the circulating ACTH levels, indicative of HPA axis activation (Feldman et al., 1981), in anesthetized rats. For this experiment, we recorded 13 CRH_PVN_ neurons (among 18 neurons recorded for the baseline firing analysis described above) that remained stable until the end of nerve stimulation session (see *Experimental Design* in Materials & Methods). Consistent with the earlier “blinded” recordings (Day et al., 1985; Hamamura et al., 1986; Kannan et al., 1987; Saphier, 1989; Saphier and Feldman, 1985, 1991), sciatic nerve stimulation increased the firing rate (time averaged spike number) of CRH_PVN_ neurons (BL 3.063 ± 1.646 Hz vs. NS 4.900 ± 1.962 Hz, paired t-test, p = 0.0108, n = 13; **Fig. 3A, B, and** **G**). Somewhat counter-intuitively, however, this activity increase was paralleled by a striking loss of RB (BL 0.1406 ± 0.1375 Hz vs. NS 0.0474 ± 0.0410 Hz, paired t-test, p = 0.0167, n = 13, **Fig. 3C, E, and** **H**), and the firing rate increased due to SS (BL 2.702 ± 1.428 Hz vs. NS 4.782 ± 1.987 Hz, paired t-test, p = 0.0025, n = 13; **Fig. 3D, F and** **I**). This finding suggests that RB paradoxically informs a certain “low activity state” of CRH_PVN_ neurons, whereas the “high activity state”, induced by stress, is due to SS at high rates.

**Figure 3.**
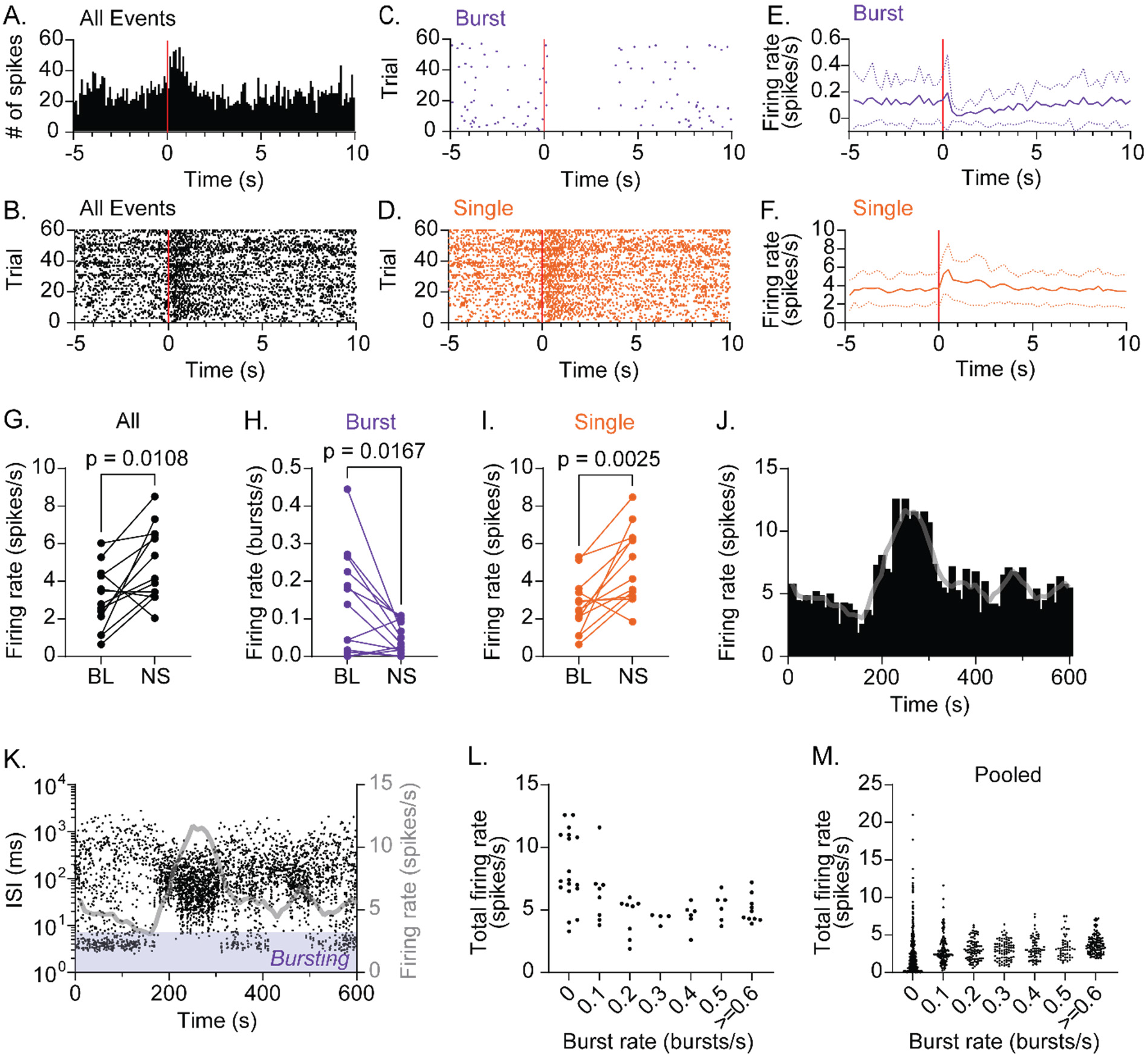
CRH_PVN_ neurons are constrained to low activity during rhythmic bursting. **A.** Peristimulus time histogram (PSTH) for a representative single-unit responding to sciatic nerve stimulation (1.5 mA, 0.5 ms x 5 pulses at 20 Hz, red line). **B-D.** Raster plots for the unit shown in A for all spikes (B), bursts (C) and single spikes (D). **E, F.** Summary time course for bursts (E) and single spikes (F). n = 13. **G-I.** Summary changes in all spikes (G, paired t-test, n = 13, p = 0.0108), bursts (H, paired t-test, n = 13, p = 0.0167) and single spikes (I, paired t-test, n = 13, p = 0.0025) before (baseline, BL) and after sciatic nerve stimulation (NS). n = 13. **J, K.** Time course of firing rate (J) and inter-spike interval (ISI) (K) for a representative single-unit during baseline recording. Grey lines are the running average (30 s) of firing rate. **L.** The relationship between burst rate and total firing rate for the representative single-unit shown in J and K. For each time bin (10 s), total spike rate was plotted against burst rate. **M.** Pooled data for all units (n = 18).

Under baseline conditions, CRH_PVN_ neurons showed spontaneous low-level firing activities; an observation in line with recent *in vivo* two-photon Ca^2+^ imaging of CRH neurons in zebra fish larvae (Berg-Maurer et al., 2016). We also found that some of CRH_PVN_ neurons show time-dependent fluctuations in their firing rate with spontaneous and transient increase (**Fig. 3J**). Thus, we next asked whether spontaneous “high activity state” is also due to an increase in SS paralleled by a loss of RB. **Figure 3J** plots the running average firing rate (10-second bins) of one representative neuron showing spontaneous emergence of high activity states. We next overlaid the time course of individual spike’s ISI to visualize the temporal relationship between the overall activity states (*i.e.* firing rate) and specific firing patterns (*i.e.* RB *vs.* SS). Similar to the stress-induced activity increase, a loss of RB temporally correlated with the emergence of the high activity state (**Fig. 3K**). On the other hand, when CRH_PVN_ neurons fired mainly with RB, their overall firing rate remained at relatively low levels. Intuitively, this is because of the brevity of burst episodes with mostly silent IBIs (**Fig. 2A, E**), precluding the high rate of SS. **Figure 3L and** **M** plot the relationship between burst and total spike rate for every 10-second time bin for this unit, as well as for all units pooled (n = 18), respectively. This analysis revealed that high levels of total spike rate only emerged when burst rate was at or near zero. Thus, our data suggest that a loss of RB permits CRH_PVN_ neurons to increase firing rate with high rate of SS. In other words, the network activity state that drives RB constrains the activity of CRH_PVN_ neurons at low levels.

### Prolonged silent periods precede burst firing

Given that the RB firing reflects a specific, low-activity state of CRH_PVN_ neurons, a question emerged: How do CRH_PVN_ neurons fire in these distinct, short burst trains? To address this, we examined the temporal properties of the burst spike trains by adopting the analysis used by Harris et al. (2001). **Figure 4A** illustrates the relationship between ISIs preceding and proceeding individual spikes of the same representative unit shown in **Figures 2A and** **B**. A cluster of spikes in the lower-left correspond with the spikes in the middle of bursts (*i.e.* both preceding and proceeding ISI are short). On the other hand, the lower-right cluster represents the initial spike of bursts (long preceding ISI and short proceeding ISI). Notably, the lack of spikes between these two clusters along the X-axis indicate that bursts preferentially start after a long silence period (*i.e.* preceding ISI >500 ms). By contrast, spikes with short preceding ISI on the left spread along the Y-axis, indicating that the end of bursts are followed by proceeding ISI of variable durations (*i.e.* not followed by an abrupt start of post-burst silence). To express these features more explicitly, we plotted the probability of burst-range firing (*i.e.* proceeding ISI <6 ms) against the preceding ISI for individual units, and then averaged for all bursting units (**Fig. 4B**). The graph shows that a burst train seldom initiates without a preceding silent period (*i.e.* ISI) of at least 200 ms, and that the likelihood of burst firing substantially increases when preceded by a silence period longer than 500 ms. In **Figure 4C**, which plots the probability of burst-range firing (*i.e.* preceding ISI <20 ms) against proceeding ISI, there was a higher probability of bursting with longer proceeding silences, but the relationship was less prominent than the “preceding ISI analysis”: this likely reflects that burst trains end with gradual prolongation of ISI (i.e. frequency adaptation). We also found that burst firing, regardless of its length, was accompanied by longer preceding silences than SS (0.33 ± 0.16 s, 0.80 ± 0.31 s, 0.89 ± 0.32 s, 1.04 ± 0.43 s, for burst lengths of single, 2, 3, and 4+, respectively; One-way ANOVA, p < 0.0001; Tukey’s multiple comparisons test, p < 0.0001, p = 0.0001, p = 0.0004 for single vs 2, 3, 4+ respectively. **Fig. 4D**). On the other hand, we did not observe these predictive trends in the “silence” following SS or bursts. Furthermore, the bursts tended to have longer proceeding silences (0.59 ± 0.36 s, 0.57 ± 0.33 s, 0.51 ± 0.40 s, for burst lengths of 2, 3, and 4+, respectively) than SS (0.37 ± 0.18 s; One-way ANOVA, p = 0.0131; Tukey’s multiple comparisons test, p = 0.0573, p = 0.0437, p = 0.2885 for single vs 2, 3, 4+ respectively. **Fig 4E**), which resulted in significant difference only between SS and bursts with 3 spikes. Again, these data reflect the fact that bursts end with gradual prolongation of ISI and are not followed by an abrupt start of a silent period. These characteristics of burst generation suggest that a prolonged silent period is followed by a brief high-frequency spike trains, which in turn are followed by a prolonged silent period, leading to RB.

**Figure 4.**
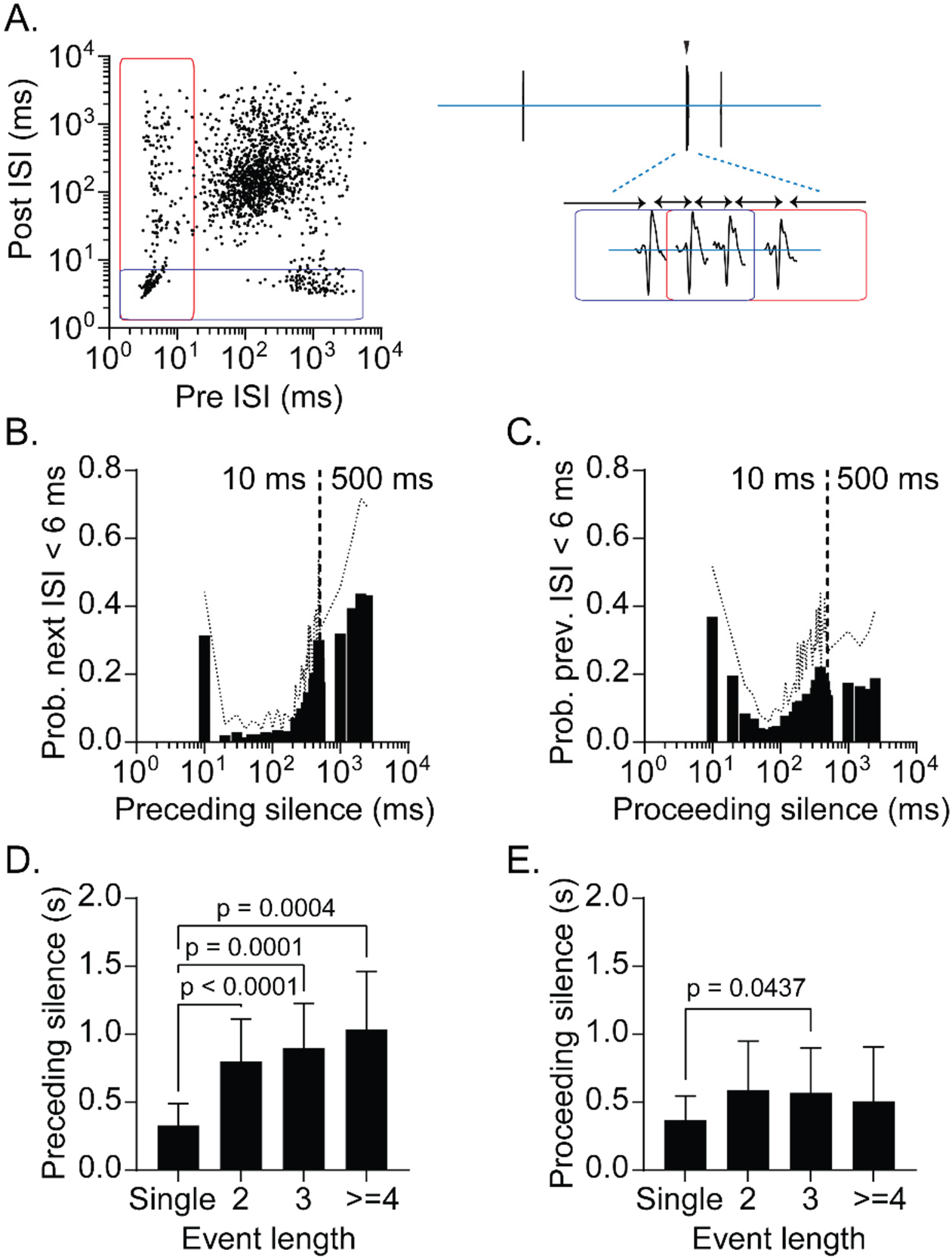
Prolonged silent periods precede burst firing. **A.** Preceding and proceeding inter-spike interval (ISI) plotted for individual spikes recorded from a representative single-unit. Blue rectangle indicates spikes with proceeding ISI <6 ms. Red rectangle indicates spikes with preceding ISI <20 ms. Left. An example of a burst episode. **B.** Summary of probability of burst firing relative to preceding silence (ISI). **C.** Summary of probability of burst firing relative to proceeding silence (ISI). **D.** Mean preceding silence as a function of event length. One-way ANOVA, p < 0.0001; Tukey’s multiple comparisons test, single vs. 2 (p < 0.0001), 3 (p = 0.0001), >=4 (p = 0.0004) **E.** Mean proceeding silence as a function of event length. One-way ANOVA, p = 0.0131; Tukey’s multiple comparisons test, single vs. 2 (p = 0.0573), 3 (p = 0.0437), >=4 (p = 0.2885) Standard deviation is represented in the graphs as dotted lines (B, C) and the error bars (D, E).

### Recurrent inhibitory circuits underlie burst firing

What are the mechanisms generating RB in CRH_PVN_ neurons? Burst firing is known to rely on the intricate interactions between intrinsic neuronal properties, synaptic inputs and network characteristics (Krahe and Gabbiani, 2004). Indeed, when studied *ex vivo* in slice recordings, in which network inputs are substantially lost, CRH_PVN_ neurons do not show overt bursting properties (Bittar et al., 2019; Jiang et al., 2019; Khan et al., 2011; Luther et al., 2002; Matovic et al., 2020; Sarkar et al., 2011; Wamsteeker Cusulin et al., 2013). In the present study, we confirmed this lack of *ex vivo* burst firing phenotype using whole-cell patch-clamp recordings in acute slices. As shown in **Figure 5A** (red traces), in response to depolarizing (monotonic) current steps, CRH_PVN_ neurons fired with a gradual increase in frequency with a moderate adaptation and they did not generate *in vivo*-like high-frequency bursts. When considering more complex network inputs that dynamically interact with the intrinsic properties of CRH_PVN_ neurons and enable RB firing, our *in vivo* burst firing pattern predicts the following: (1) the intrinsic properties of CRH_PVN_ neurons favour high-frequency burst firing after a prolonged silence period, and (2) network inputs that cause the prolonged silence contribute to the slow “rhythms’’ of bursting. Taken together, these features are indicative of an inhibitory-driven recurrent network burst mechanism (McCormick and Feeser, 1990; Sherman, 2001; Steriade et al., 1993). Supporting this idea, early *in vivo* recordings of PVN neurons predicted the existence of recurrent inhibitory circuits based on inhibition following antidromically-elicited action potentials (Saphier and Feldman, 1985). More recent studies identified a group of local GABAergic interneurons that form recurrent inhibitory circuits (Jiang et al., 2018, 2019; Ramot et al., 2017). Thus, we hypothesized that recurrent inhibitory circuits underlie the RB of CRH_PVN_ neurons. To test this, we computationally studied the dynamics of single CRH_PVN_ neurons within a network by using a spiking network of adaptive-exponential (AdEx) neuron models (Brette and Gerstner, 2005; Gerstner and Naud, 2009; Izhikevich, 2003; Markram et al., 2015). First, the single neuron AdEx models (Brette and Gerstner, 2005) were fit to slice patch-clamp recordings of CRH_PVN_ neurons (**Fig. 5A and Table 1**). We evaluated the performance of the model by quantifying the mean squared difference in the subthreshold traces (**Fig. 5A**) and the frequency-current (F-I) curve (**Fig. 5B**) between the model and experimental data (Brette and Gerstner, 2005). Our simulations show that the single neuron models can capture a rapid spike adaptation that follows the first few spikes, timing of repetitive spike firing and the F-I relationship for a series of square pulse depolarizing current injections (**Fig. 5A and** **B**).

**Figure 5.**
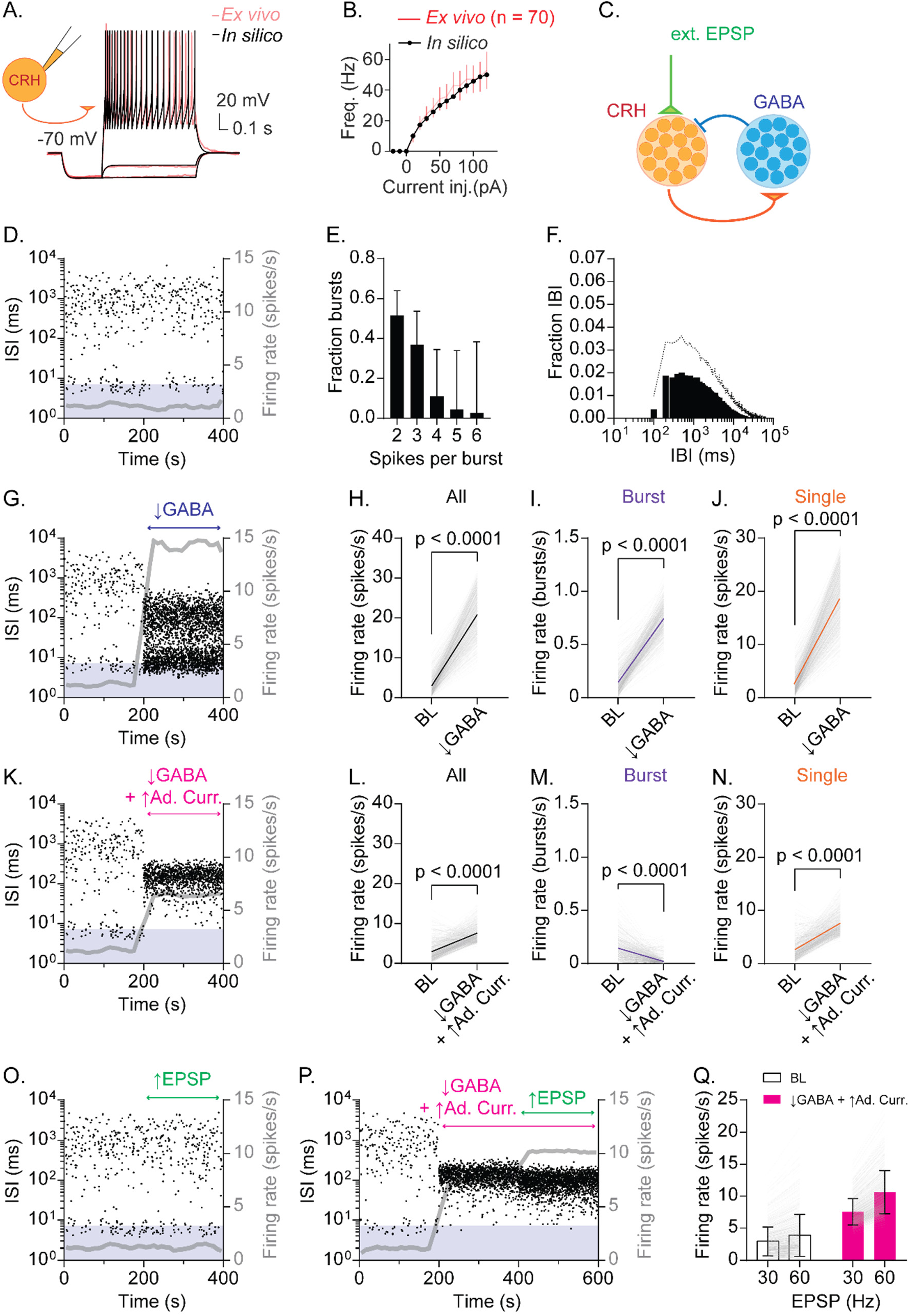
Recurrent inhibitory circuits generate burst firing and gate firing response to excitatory inputs in CRH_PVN_ neurons. **A.** CRH model neuron *in silico* (black) fitted to current-clamp recordings of CRH_PVN_ neurons *ex vivo* (red). **B.** F-I curves for *in silico* (black) and *ex vivo* (red) neurons. **C.** Network model diagram. **D.** Time course of inter-spike interval (ISI) for a representative model CRH neuron. Gray line is the running average firing rate. **E.** Summary burst length distribution for model simulations (n = 500). **F.** Summary IBI distribution for model simulations (n = 500). **G.** An example ISI time course before and after a drop of GABA release (Pr 0.8 → 0.1) from 200 s. Gray line is the running average firing rate. **H-J.** Summary changes in all spikes (H, paired t-test, p < 0.0001, n = 500), bursts (I, paired t-test, p < 0.0001, n = 500) and single spikes (J, paired t-test, p < 0.0001, n = 500) before (BL) and after the GABA release drop. **K.** An example ISI time course before and after a drop of GABA release (Pr 0.8 → 0.1) combined with an increase in spike-triggered adaptation current from 200 s. Gray line is the running average firing rate. **L-N.** Summary changes in all spikes (L, paired t-test, p < 0.0001, n = 500), bursts (M, paired t-test, p < 0.0001, n = 500) and single spikes (N, paired t-test, p < 0.0001, n = 500) before (BL) and after combined removal of GABA release and increased spike-triggered adaptation current. **O.** An example ISI time course before and after EPSP frequency increase (30 Hz → 60 Hz) from 200 s. Gray line is the running average firing rate. **P.** An example ISI time course before and after the combined change in GABA release and spike-triggered adaptation current, followed by an increase in EPSP frequency (30 Hz → 60 Hz) from 400 s. **Q.** Summary graph for changes in firing rate before (30 Hz) and after EPSP frequency increase (60 Hz) with (white) and without recurrent inhibition (pink). Two-way ANOVA (EPSP x Recurrent inhibition interaction, p < 0.0001; Tukey’s multiple comparisons test, BL-30 Hz 2.942 ± 2.220 Hz vs. BL-60 Hz 3.895 ± 3.252 Hz, p < 0.0001; ↓GABA+↑Ad. Curr.-30 Hz 7.548 ± 2.068 Hz vs. ↓GABA+↑Ad. Curr.-60 Hz 10.630 ± 3.387 Hz, p < 0.0001). Standard deviation is represented in the graphs as the error bars.

**Table 1:**
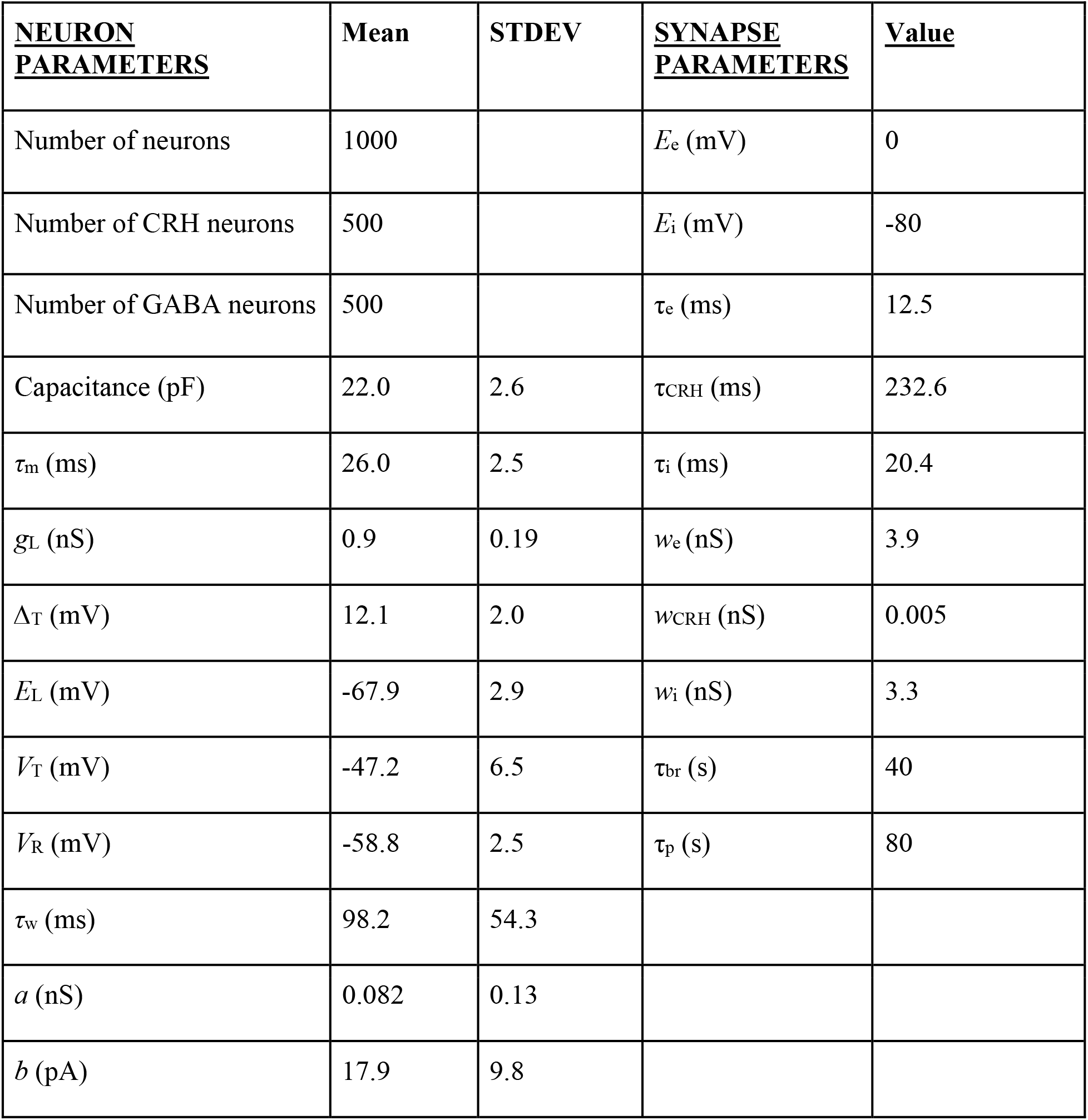

| <u>NEURON</u><br><u>PARAMETERS</u> | Mean | STDEV | <u>SYNAPSE</u><br><u>PARAMETERS</u> | <u>Value</u> |
| --- | --- | --- | --- | --- |
| Number of neurons | 1000 | | $E_e$ (mV) | 0 |
| Number of CRH neurons | 500 | | $E_i$ (mV) | -80 |
| Number of GABA neurons | 500 | | $\tau_e$ (ms) | 12.5 |
| Capacitance (pF) | 22.0 | 2.6 | $\tau_{CRH}$ (ms) | 232.6 |
| $\tau_m$ (ms) | 26.0 | 2.5 | $\tau_i$ (ms) | 20.4 |
| $g_L$ (nS) | 0.9 | 0.19 | $w_e$ (nS) | 3.9 |
| $\Delta_T$ (mV) | 12.1 | 2.0 | $w_{CRH}$ (nS) | 0.005 |
| $E_L$ (mV) | -67.9 | 2.9 | $w_i$ (nS) | 3.3 |
| $V_T$ (mV) | -47.2 | 6.5 | $\tau_{br}$ (s) | 40 |
| $V_R$ (mV) | -58.8 | 2.5 | $\tau_p$ (s) | 80 |
| $\tau_w$ (ms) | 98.2 | 54.3 | | |
| $a$ (nS) | 0.082 | 0.13 | | |
| $b$ (pA) | 17.9 | 9.8 | | |

Next, the fitted single-neuron models were integrated into a spiking network model to test whether the CRH-GABA neuron network generates rhythmic bursting. We have constructed a network model of the PVN that contains 500 CRH neurons and 500 GABA neurons (**Fig. 5C**). The excitatory CRH neurons send a sparse projection to the inhibitory GABA population (2% connection probability), and the GABA population projects back to the CRH neurons with a similar, sparse projection (2%) (Destexhe, 2009). CRH neurons also receive noisy random synaptic inputs (30 Hz fast EPSPs), approximated by an instantaneous rise and exponential decay (Rothman and Silver, 2014). GABAergic neurons, in turn, receive excitatory CRHergic inputs that have slow rise and decay time constant (∼200ms), modelled as single time constant alpha synapse (Destexhe et al., 1994), reflecting the slow CRHR1 mediated excitation (Jiang et al., 2019; Ramot et al., 2017). The putative network construction was then optimized to fit our *in vivo* observations. In brief, from a representative unit (shown in **Fig. 3K**), temporal subsections were isolated as representative RB and SS firing patterns. Then, in the network model, several rounds of a genetic algorithm were used to optimize the synaptic network parameters; simulations were evaluated using the earth movers’ distance (EMD) between the log ISI distributions of the representative spike train and simulated spike trains (Materials and Methods). In this optimized network setting, we found that model CRH neurons generate RB (**Fig. 5D**). The model CRH neurons, which have heterogeneity in their intrinsic properties obtained in *ex vivo* recordings (**Table 1**), showed heterogeneity in their burst rates (0.1456 ± 0.1072 bursts/s, n = 500). By further characterizing burst firing properties of these bursting model CRH neurons, we found that our network model recapitulated several *in vivo* features, including burst length and IBI (**Fig. 5E, F**). Consistent with these features, RB in the model also constrained the overall firing rate at low levels (∼2 Hz, **Fig. 5D gray line**). These data show that CRH_PVN_ neurons (which are not intrinsically bursting) fire in RB within a recurrent inhibitory network.

Next, to test the causal roles of recurrent inhibition in constraining firing rate, we performed a series of tests with the network. First, a reduction of recurrent inhibition (by stochastically lowering the release probability (Pr) of GABA→CRH synapses from [0.9-1.0] to [0.01-0.2]) alone with no additional change in external excitatory inputs, robustly increased spiking, and consequently the overall firing rate (**Fig. 5G-J**). This indicates that recurrent inhibition constrains the overall firing activity of CRH neurons. However, the removal of recurrent inhibition alone did not eliminate the burst-range high-frequency firing in CRH neurons (**Fig. 5G, L**). Interestingly, we found that the *in vivo*-like switch from RB to SS can be achieved by an additional parameter change, an increase in the “spike-triggered adaptation current” from [5-18] pA to [36-50] pA (Brette and Gerstner, 2005) (**Fig. 5K-N**). This result predicts potential intrinsic single-cell properties underlying the generation of RB (or its prevention). For example, the spike-triggered adaptation current can represent after-hyperpolarization, (a loss of) after-depolarization, or both (Izhikevich, 2003). Next, our modeling data also indicate that recurrent inhibition constrains the response (spike outputs) of CRH neurons to excitatory inputs. To test this idea directly, we increased external synaptic inputs under RB (*i.e.* in the presence of recurrent inhibition) and SS (*i.e.* in the absence of recurrent inhibition plus an increase in spike-triggered adaptation current) states (**Fig. 6 O, P**). In the presence of recurrent inhibition, an increase in excitatory synaptic inputs (from 30 Hz to 60 Hz) caused small change in the overall firing rate, namely CRH neurons’ response (**Fig. 5O, Q**). By contrast, in the absence of recurrent inhibition, an identical increase in excitatory synaptic inputs robustly increased the firing rate of CRH neurons (**Fig. 5P, Q**, Two-way repeated ANOVA, EPSP x recurrent inhibition, interaction p < 0.0001), showing that recurrent inhibition works as a gain regulator at CRH neurons.

**Figure 6.**
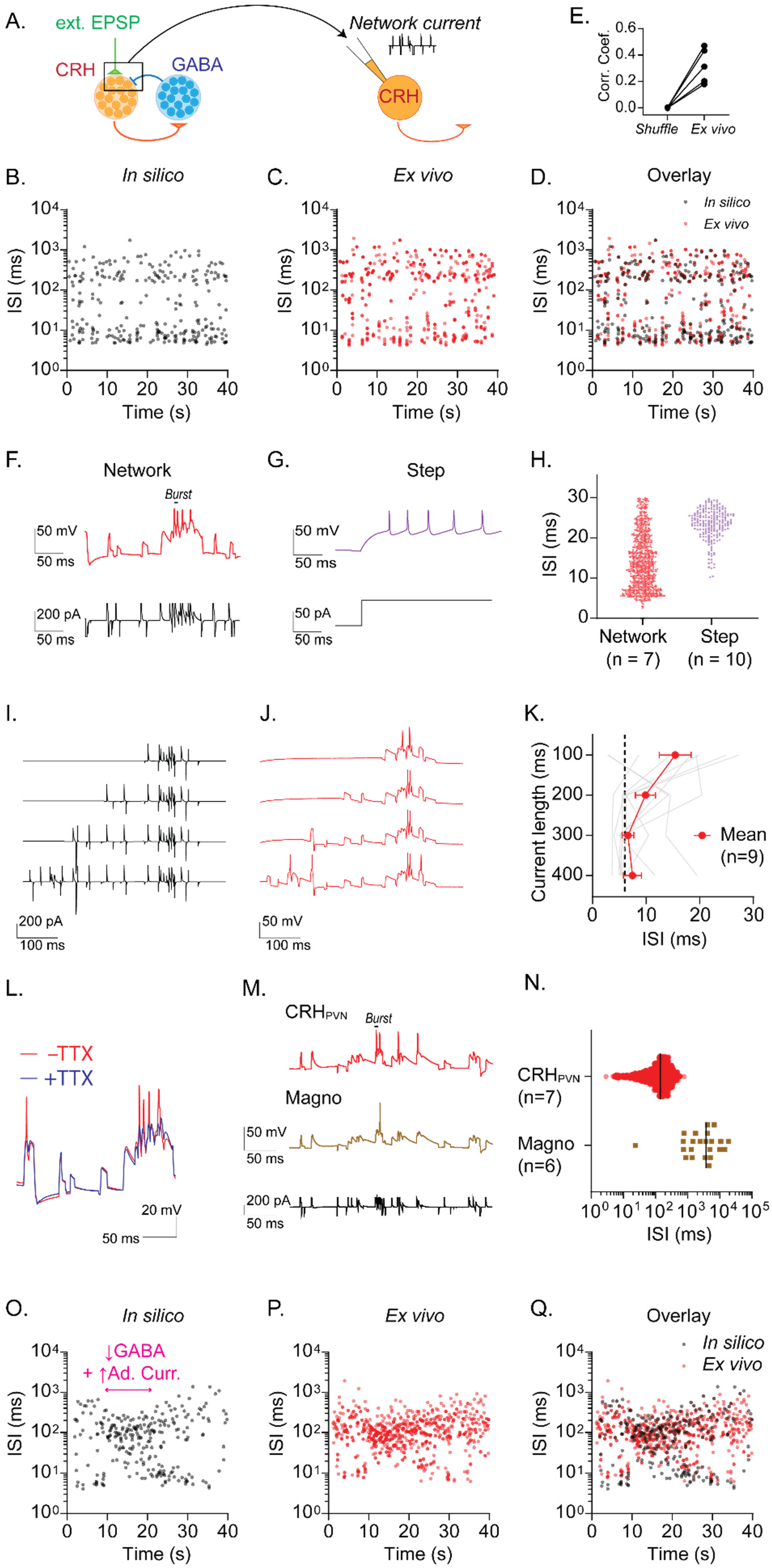
Recurrent inhibitory circuits underlie burst firing. **A.** Diagram of “network clamp” experiment. **B-D.** Time course of inter-spike interval (ISI) for a representative model CRH neurons (B), a biological CRH_PVN_ neuron injected with network current (C) and overlay (D). **E.** Correlation coefficient between the time series of model neuron spike train *in silico*, a biological CRH_PVN_ neurons’s spike train *ex vivo*, and a shuffled spike train consisting of the model spike train with ISI’s shuffled (n = 6). **F, G.** Spike firing patterns in response to network current (F) and depolarizing current step (G) in a biological CRH_PVN_ neuron. **H.** ISI distribution of spikes generated by network current (red) and steps of current (purple). Note that spikes with ISI smaller than 30 ms were used for comparison. **I, J.** Network currents of variable length (I) and corresponding membrane voltage changes in CRH_PVN_ neurons (J). **K.** Summary graph for the shortest ISI triggered by network currents of variable lengths. **L**. CRH_PVN_ neuron’s firing response to network current with (blue) and without tetrodotoxin (TTX, 1 µM, red). **M.** Firing of CRH_PVN_ neuron (orange) and magnocellular neuron (brown) in response to an identical network current (black). **N.** Summary of ISI triggered by network current in CRH_PVN_ and magnocellular neurons. **O-Q.** A representative ISI time course for the transition between rhythmic burst and single spiking in a model CRH neuron (O), a biological CRH_PVN_ neuron injected with the network current of the model neuron shown in O, and overlay (Q).

### Recurrent inhibition generates burst firing in single CRH_PVN_ neurons

Our model simulation predicted that recurrent inhibition is sufficient and necessary for RB and constrains the firing rate of CRH_PVN_ neurons. However, it remains unknown whether our single neuron model, while grounded in direct physiological measurements (**Fig. 5AB**, **Table 1**), captures essential biophysical properties required for *in vivo*-like firing patterns. To directly test this, we went back to *ex vivo* slice patch clamp electrophysiology and asked if an injection of whole-cell (somatic) currents that mimic the recurrent network inputs of the spiking network model is sufficient to drive RB firing in biological CRH_PVN_ neurons (**Fig. 6A**). To this end, we first selected a model CRH_PVN_ neuron that showed representative RB firing (**Fig. 6B**), and then extracted the “network currents” that this model neuron receives within the network model (see Materials and Methods). We then injected these “network currents” into biological CRH_PVN_ neurons in slices using current-clamp electrophysiology (**Fig. 6C**). We found a remarkable similarity between the RB firing patterns of a model neuron and a biological CRH_PVN_ neuron that received the “network currents” (**Fig. 6D, E**). Notably, in response to square pulse depolarization, biological CRH_PVN_ neurons in acute slices fire in regular spiking patterns (**Fig. 6F, H**) with the lowest ISI >10 ms. The same neurons, however, readily generated burst range high-frequency firing (ISI <6 ms) in response to complex network current injections derived from the network model (**Fig. 6G, H**). A closer examination of the network inputs indicated that high-frequency excitatory inputs drove burst firing (**Fig. 6G**). Importantly, however, the high-frequency inputs alone were not sufficient to elicit burst firing, and preceding network inputs were also important. **Fig. 6I-K** show that truncated current injection only containing the high-frequency inputs failed to elicit burst firing, and that preceding noisy network inputs (which caused subthreshold membrane potential fluctuations) were necessary to drive the burst firing. These results corroborate with *in vivo* data that burst firing almost exclusively occurred after a prolonged (>200 ms) silent period (**Fig. 4C**), indicating the importance of dynamic interactions between network synaptic inputs (that generate subthreshold membrane potential changes) and intrinsic properties. The burst firing triggered by network synaptic currents were abolished in the presence of voltage-gated Na^+^ channel blocker tetrodotoxin (TTX, 1 µM, **Fig. 6L**), confirming that they were indeed action potentials. Moreover, we found that in magnocellular (non-CRH) neuroendocrine neurons of the PVN, which are known to have *ex vivo* intrinsic properties (Luther et al., 2002) and *in vivo* firing patterns (Leng and MacGregor, 2018) different from CRH_PVN_ neurons, the identical network current failed to trigger the RB (**Fig. 6M, N**). Lastly, **Figure 6O** shows an example of model neurons capturing *in vivo*-like alternations of RB and SS using a transient change in the recurrent inhibition and adaptation current (see *in vivo* data in **Fig. 4J**). The injection of the network currents effectively elicited a similar transition of firing pattern in biological CRH_PVN_ neurons in *ex vivo* (**Fig. 6P, Q**).

## DISCUSSION

Here, we report single-unit activities of identified CRH_PVN_ neurons *in vivo* for the first time. We show that CRH_PVN_ neurons, within intact circuits, can fire distinct brief bursts, a firing pattern that has not been observed in *ex vivo* slice electrophysiology (Bittar et al., 2019; Jiang et al., 2019; Khan et al., 2011; Luther et al., 2002; Matovic et al., 2020; Sarkar et al., 2011; Wamsteeker Cusulin et al., 2013). Guided by the *in vivo* firing patterns, we have developed a computational model and showed that the RB firing mode reflects critical roles of recurrent inhibition in constraining the overall firing rate of CRH neurons. More generally, the recurrent inhibition controls the gain of CRH neurons’ response to excitatory inputs. In biological CRH_PVN_ neurons *ex vivo*, the injection of whole-cell currents derived from our network model effectively triggered the *in vivo*-like RB and recapitulated the transition from RB to SS, thus providing direct evidence that intrinsic properties of CRH_PVN_ neurons, which do not overtly fire in burst in slices, are capable of firing *in vivo*-like RB by interacting with complex inputs reflecting network characteristics. In summary, using a combination of experimental and computational approaches, we demonstrate a novel circuit mechanism that controls state-dependent activity switch of CRH_PVN_ neurons between the baseline and stress.

### CRH_PVN_ neurons fire brief bursts with long inter-burst intervals

Pioneering studies in 1980s used antidromic activation of the median eminence and reported single-unit recordings from parvocellular neuroendocrine neurons of the PVN, a class of neurons that include CRH and several other hormone-releasing neurons, in anesthetized rats (Day et al., 1985; Hamamura et al., 1986; Kannan et al., 1987; Saphier, 1989; Saphier and Feldman, 1985). Here, using the optrode technique (Lima et al., 2009), we report firing activities of identified CRH_PVN_ neurons in anesthetized mice. Overall, our results from CRH_PVN_ neurons agree with the early results from parvocellular neurons in that both populations show spontaneous firing at low rates, and that majority of them increased firing rates in response to stress-mimicking peripheral nerve stimulation.

A key new finding of our study is that CRH_PVN_ neurons fire distinct bursts characterized by RB, a brief train of high-frequency action potentials (∼ 3 action potentials/bursts at >100 Hz) intervened between long periods of silence (constraining the overall firing rate), in addition to SS with more variable and lower spike-to-spike frequencies. This RB firing was not explicitly documented by the earlier studies in rats (Day et al., 1985; Hamamura et al., 1986; Kannan et al., 1987; Saphier, 1989; Saphier and Feldman, 1985; Watanabe et al., 2004); likely because these studies analyzed the firing activities only as time-binned firing rates and did not examine spike-to-spike temporal patterns. In fact, representative traces in one of aforementioned studies showed spikes >100 Hz (see Fig. 4 of (Saphier and Feldman, 1985)). It should be emphasized that the RB we report here is distinct from “phasic (burst) firing” which is widely known as a signature firing patterns of vasopressinergic magnocellular neurons of the PVN and SON (Brown and Bourque, 2006). The magnocellular phasic burst consists of lower intra-burst frequencies (2.5-20 Hz) and lasts much longer with continuous firing (>10 s) than the RB (<100 ms) we report here in CRH_PVN_ neurons (Poulain and Wakerley, 1982; Watanabe et al., 2004). Also, the phasic burst firing is followed by a prolonged (∼10 s) silent period generated by intrinsic mechanisms (Brown and Bourque, 2006). We did not observe this type of phasic bursts in identified CRH_PVN_ neurons consistent with the fact magnocellular neurons do not express CRH (Biag et al., 2012; Simmons and Swanson, 2009).

The brief bursts were intervened with mostly silent IBIs of its median length around 2 s during the RB state. Consequently, despite the high frequency (>100 Hz) burst firing, the time-averaged firing rate of the RB state remained low (3-5 Hz). Further, during non-stress conditions, the RB state dynamically shifted to a non-bursting, SS state where CRH_PVN_ neurons increased their firing rate (10-20 Hz) due to continuous single spiking. A similar shift from RB to continuous SS was evident when CRH_PVN_ neurons increase their firing rate in response to stress-mimicking sciatic nerve stimulations. When considering hormonal CRH levels, the drive for the hormone release is likely to be coded by firing rate over the time scale of seconds and longer, rather than millisecond precisions of firing patterns. Thus, we speculate that persistent SS firing and ensuring elevation of spike rate play a major role in the hormonal CRH release. On the other hand, it raises an intriguing possibility that RB firing may encode and convey specific information to specific downstream targets. This topic warrants future investigation in light of emerging new roles of CRH_PVN_ neurons beyond hormonal release of CRH, including wakefulness (Ono et al., 2020), valence encoding (Kim et al., 2019a), reward processing (Yuan et al., 2019), and defensive behavior control (Daviu et al., 2020; Füzesi et al., 2016) under non-stress and stress conditions.

### A computational model predicts a novel disinhibition mechanism of CRH_PVN_ neurons

Decades of research has established that, under no-stress conditions, the excitability of CRH_PVN_ neurons is tonically constrained by powerful GABAergic synaptic inhibition, and that a release from this tonic inhibition, or disinhibition, is one dominant mechanism for the activation of these neurons, and consequently the HPA axis (Bains et al., 2015; Cole and Sawchenko, 2002; Cullinan et al., 2008; Inoue and Bains, 2014, 2014; Levy and Tasker, 2012; Roland and Sawchenko, 1993; Ulrich-Lai and Herman, 2009). Multiple disinhibitory mechanisms have been identified, likely reflecting diverse and overlapping neural and hormonal pathways controlling the activities of the HPA axis (Joëls and Baram, 2009; Ulrich-Lai and Herman, 2009). These include the inhibition of the presynaptic GABAergic neurons (Anthony et al., 2014; Johnson et al., 2019), depression of GABAergic synaptic terminals (Ferri and Ferguson, 2005; Han et al., 2002; Hewitt and Bains, 2006; Khazaeipool et al., 2018) and a depolarizing shift in the equilibrium potential of GABA_A_ receptor in the postsynaptic CRH_PVN_ neurons (Hewitt et al., 2009; Sarkar et al., 2011). However, it remained unknown how GABAergic inhibition controls the firing activities of CRH_PVN_ neurons *in vivo*. In this study, we developed a simple computational model, which was guided by distinct brief burst firing *in vivo*, and revealed previously underappreciated contribution of recurrent inhibition to the tonic inhibition of CRH_PVN_ neurons. Specifically, the feedback inhibition effectively prevented continuous firing and created a lasting (∼2 s) silent periods in between brief bursts, and consequently constrained the time-averaged firing rate low (∼3 Hz). Strikingly, a decrease of recurrent inhibition, with no additional change in the excitatory inputs, profoundly increased the overall firing rate of CRH_PVN_ neurons. In other words, (a loss of) the recurrent inhibition alone partially recapitulates the disinhibitory mechanisms of CRH_PVN_ neurons. That said, there is more to consider regarding the activity of CRH_PVN_ neurons, as the removal of recurrent inhibition alone results in continuous burst-range high frequency firing, which does not typically occur during stress-induced firing increase *in vivo* (**Fig. 3A-F**). Thus, our model predicts that the *in vivo*-like transition from RB to SS requires an increase in spike-triggered adaptation current (Brette and Gerstner, 2005), pointing to single-neuron intrinsic properties underlying burst generation. For example, the spike-triggered adaptation current can represent after-hyperpolarization, (a loss of) after-depolarization, or both (Brette and Gerstner, 2005).

It should be noted that our model for the recurrent excitatory-inhibitory circuit in the PVN is simplified and does not incorporate external inhibitory inputs to CRH_PVN_ neurons or external excitatory and inhibitory inputs to the GABAergic neurons (Ulrich-Lai and Herman, 2009). However, this model parsimoniously captures the key features of firing behavior *in vivo*, including the brevity of burst episodes and long IBI (**Fig. 5E, F**). Thus, our simplified model, which is the first network model for CRH_PVN_ neurons to our best knowledge, is a useful first step to understand the spiking behaviors of CRH_PVN_ neuron in a network, and to generate experimentally testable hypotheses for future research.

So, what are the identities of the GABAergic neurons that form recurrent connectivity with CRH_PVN_ neurons? Indeed, the recurrent inhibitory circuits for CRH_PVN_ neurons has been first suggested by an early *in vivo* electrophysiology recordings of parvocellular neurons in rats based on an inhibition following action potentials antidromically activated from the median eminence (Saphier and Feldman, 1985). Recent studies have revealed local recurrent circuits within the PVN, where CRH_PVN_ neurons excite CRHR1-expressing GABAergic interneurons, which in turn send GABAergic inputs to CRH_PVN_ neurons (Jiang et al., 2018). CRH_PVN_ neurons excite CRHR1-expressing neurons via slow metabotropic actions with little, if any, fast (glutamatergic) excitatory synaptic transmission (Jiang et al., 2018; Ramot et al., 2017). Notably, our network model found that the long IBI (∼2 s) of brief bursts require slow excitatory EPSPs consistent with metabotropic signaling (τ_CRH_ >200 ms) rather than fact ionotropic EPSPs (τ_e_ <20 ms). In addition to this intra-PVN microcircuit, it is possible that CRH_PVN_ neurons form additional recurrent inhibitory circuits. Indeed, there is growing appreciation that CRH_PVN_ neurons send projections within the brain besides their well established projections to the median eminence (for hormonal CRH release) (Füzesi et al., 2016; Jiang et al., 2018, 2019; Kim et al., 2019a; Li et al., 2020; Ono et al., 2020; Ramot et al., 2017; Rho and Swanson, 1989; Yuan et al., 2019). For example, the peri-fornical area (Füzesi et al., 2016; Rho and Swanson, 1989) and the lateral hypothalamus (Li et al., 2020; Ono et al., 2020) receive direct synaptic inputs from CRH_PVN_ neurons: these brain areas in turn send direct GABAergic inputs to CRH_PVN_ neurons (Boudaba et al., 1996; Cullinan et al., 2008; Roland and Sawchenko, 1993; Ulrich-Lai and Herman, 2009).

### Synaptic activity underlies flexible firing patterns of CRH_PVN_ neurons *in vivo*

In acute slices *ex vivo*, CRH_PVN_ neurons typically show a SS electrophysiological phenotype, or “regular” spiking, with up to around 50 Hz of spike-to-spike frequency in response to steps of intracellular current injections in mice and rats (Bittar et al., 2019; Jiang et al., 2019; Khan et al., 2011; Luther et al., 2002; Matovic et al., 2020; Sarkar et al., 2011; Wamsteeker Cusulin et al., 2013). Thus, our *in vivo* recordings of the RB (>100 Hz spike-to-spike frequency) revealed a previously underappreciated repertoire of the firing patterns of CRH_PVN_ neurons. In other words, our data suggest that the intrinsic properties of CRH_PVN_ neurons conduct flexible input/output signal processing by interacting with the physiological network dynamics which is substantially lost in *ex vivo* slice preparations. Supporting this idea, our modeling showed that model CRH neurons, which is fitted to “regular” spiking phenotype characterized *ex vivo*, readily fire in RB when they receive the bombardment of excitatory inputs (30 Hz EPSPs) combined with the recurrent inhibition. Thus, our modeling results demonstrate that as simple as two types of network-like inputs are sufficient to generate a firing pattern that is qualitatively different from and not readily evident in conventional (*e.g.* square pulse) electrophysiological characterizations *ex vivo*.

The flip side of our modeling approach is the oversimplification of the network inputs and questions about their physiological relevance. To directly address the latter concern, we applied the modeling results to biological neurons *ex vivo*. Specifically, we extracted network inputs from representative model neurons that fire RB, and then injected the network current waveform into biological neurons under whole-cell current-clamp configuration *ex vivo*. We term this new approach as “network clamp”. Network clamp injects a predetermined current waveform which can be readily available from the leaky-integrate-and-fire model, and therefore much simpler, operationally and computationally, than dynamic clamp. One limitation is that the amplitude of the current fluctuations can be larger than physiologically realistic range. Despite this limitation, however, we found that the network current generated RB in CRH_PVN_ neurons and the transition to SS upon changes in network parameters (*i.e.* removal of recurrent inhibition and increase in adaptation current), providing a direct evidence for the capability of CRH_PVN_ neurons to fire with high frequency bursts (*i.e.* RB pattern) and also bidirectionally switch to regular spiking (*i.e.* SS pattern). Importantly, the burst firing does not simply occur as a results of current injection patterns but requires unique interaction between the intrinsic properties of CRH_PVN_ neurons and the network inputs. This is supported by the following two controls. First, network inputs, which drove subthreshold membrane potential changes preceding (>200 ms) the burst firing, were necessary because truncated (<200 ms) network currents, which only contain high-frequency synaptic inputs component, failed to elicit the brief bursts in CRH_PVN_ neurons (**Fig. I-K**). Second, magnocellular neurons, a separate class of neuroendocrine neurons of the PVN with different intrinsic properties (Luther et al., 2002), did not fire in brief bursts in response to the same network current injection that elicited brief bursts in CRH_PVN_ neurons (**Fig. 6M, N**).

These results point to future applications of the “network clamp” in order to examine specific biophysical mechanisms that interact with network inputs to generate *in vivo*-like firing patterns in other brain areas. It is well known that neurons in slice differ in their response properties from intact *in vivo* preparations (Destexhe and Paré, 1999; Destexhe et al., 2003) and that injection of noise approximating the inputs from the large number of synaptic contacts in intact preparations can recreate firing conditions observed *in vivo* (Destexhe and Rudolph-Lilith, 2012; Destexhe et al., 2003; Rudolph and Destexhe, 2003, 2006). Advanced techniques have been developed to measure neuronal firing rate responses under injection of theoretically motivated noise processes (Zerlaut et al., 2016). In this work, however, we have focused on injecting the time-varying inputs from a *single* cell in the network model to a biological neuron, so that we can directly compare the activity patterns evoked in the biological and simulated cells. We find that computationally generated inputs can recreate the specific burst firing mode we have reported in this work. In this way, this injection protocol bears some similarity to the “iteratively constructed network (ICN)” developed by Alex Reyes (Reyes, 2003) to provide a direct test of whether biological cells can propagate synchrony without some of the caveats that can be imposed by computational simulations (Rudolph and Destexhe, 2007). We suggest that this “network clamp” protocol has more general applicability, however, for validating spiking network models of neural circuits. In this work, the spiking network model provided a testable prediction: initially, biological neurons recorded *ex vivo* did not exhibit bursting behavior when stimulated with step current pulses, but would they burst when driven with network-generated input? In this case, the biological neuron in this “network clamp” paradigm exhibited behavior consistent with this prediction, strengthening confidence that our spiking network model captures a meaningful underlying mechanism in this system. Because this paradigm allows to test multiple model predictions in a single setup (for conventional patch-clamp electrophysiology), and because it also opens the possibilities for testing how neuronal firing and response patterns may change with neuromodulation applied in the slice, we suggest that the model-experiment protocol developed and tested in this work may be able to distinguish between multiple models that can account for a single dataset in isolation (Marder and Goaillard, 2006).

## MATERIALS & METHODS

### Animals

All experimental procedures were performed in accordance with the Canadian Council on Animal Care guidelines and approved by the University of Western Ontario Animal Use Subcommittee (AUP: 2018-130). Homozygous *Crh-IRES-Cre (B6(Cg)-Crhtm1(cre)Zjh/J)* mice (Stock No: 012704, the Jackson Laboratory) were crossed with homozygous Cre-reporter *Ai14 (B6.Cg-Gt(ROSA)26Sortm14(CAG-TdTomato)Hze/J)* mice (Stock No: 007908, the Jackson Laboratory) to produce CRH-TdTomato reporter offspring. The specificity of cre expression in the PVN in these mice has been characterized in previous studies (Chen et al., 2015; Wamsteeker Cusulin et al., 2013). Ten adult male mice (> 60 days old) were used for viral injections and recordings. Animals were group-housed (2-4 per cage) in standard shoebox mouse cages with *ad libitum* access to food and water. Animals were housed on a 12-hour dark/12-hour light cycle (lights on from 07:00-19:00) in a temperature-controlled room (23 ± 1°C).

### Viral injection

To express channelrhodopsin-2 (ChR2) in CRH_PVN_ neurons, we used an adeno-associated virus (AAV) carrying the EF1α promoter and the double-floxed inverted ChR2(H134R)-EYFP coding sequence that is inverted and turned on by Cre-recombination (AAV2/5- EF1α-DIO- ChR2(H134R)-EYFP). The plasmid was a gift from Dr. Karl Deisseroth http://n2t.net/addgene:20298 ; RRID:Addgene_20298). The AAV preparation was obtained from the Neurophotonics Centre (5 x 10^13^ GC/ml, Laval University, Canada).

For AAV injection, the animal was anesthetized under isoflurane (2%) using a low flow gas anesthesia system, (Kent Scientific Corporation) and placed in a stereotaxic apparatus on a heating pad. A finely pulled glass capillary was loaded with the virus and slowly lowered into the brain of animals, targeting the PVN on each side of the brain (A/P: -0.70 mm, M/L: ± 0.25 mm, and D/V: -4.75 mm from bregma). A total of ∼240 nL (41.4 nL x 6 at 23 nL/sec) was pressure-injected on each side using the Nanoject II (Drummond Scientific Company). The pipette was held in place for 5 minutes after injection to allow diffusion before being slowly retracted from the brain. The incision was closed by suturing and the animals were injected with analgesic (buphrenorphone 0.1 mg/kg, s.c.) at the end of surgery. Animals were allowed to recover for at least 6 weeks for optimal ChR2 expression before electrophysiological recordings.

### Electrophysiology

Animals were initially anesthetized under isoflurane (1-2%) and urethane (1.5 g/kg in 0.9% saline, intraperitoneal) to perform surgery for sciatic nerve isolation and a craniotomy/durotomy for insertion of the recording probe into the brain. We first isolated the sciatic nerve (see details in *Sciatic nerve stimulation*), and then the probe was lowered into the brain. Thereafter, the animal was taken off isoflurane. The probe was vertically inserted above the PVN (A/P: -0.70 mm from bregma, M/L: ± 0.25 mm from bregma) and slowly lowered ventrally to the PVN (target D/V: - 4.5 mm from cortical surface, photo-tagged units (see details in *Optogenetic identification of CRH neurons*) were found between -4.20 mm to -4.80 mm from cortical surface).

Extracellular neural signals were recorded using a single shank, 32-channel (8 rows x 4 columns) silicon probe, with recording sites spanning 60 µm in depth (Cambridge NeuroTech). An optic fiber (100 µm core diameter, 0.37 NA, A45; Doric Lenses) was attached by the manufacturer parallel to the electrode shank with a vertical offset of 250 µm, constituting an optrode. The electrode was connected to a digital headstage (ZD ZIF-Clip; Tucker-Davis Technologies) with an internal Intan amplifier chip. The digitized signals were sent to the amplifier (PZ5; Tucker-Davis Technologies), which in turn connects out to the processing unit (RZ5D, Tucker-Davis Technologies) via a fiber optic cable. Broadband signals were sampled at 25 kHz and bandpass filtered between 300-3000 Hz. The threshold for spike detection was a minimum three standard deviations above the noise floor. The waveforms for detected spikes were sorted offline (Offline Sorter, Plexon) using manual and automatic clustering (T-Distribution E-M). To ensure the quality of single-unit isolation, only discrete clusters with L-ratios < 0.05 (in 2D or 3D spaces) was included for single-unit analysis.

### Optogenetic identification of CRH neurons

TTL signals to trigger optogenetics light stimulation were sent from the signal processing and acquisition software Synapse (Tucker-Davis Technologies) to a LED driver (PlexBright LD-1 Single Channel LED Driver, Plexon). The LED driver controlled the intensity of light emitted from the LED module (PlexBright Table-Top LED Module Blue 465 nm, Plexon) which was coupled to an optic patch cable (200 µm core, PlexBright Optical Patch Cable, Plexon). The patch cable connected via a plastic sleeve to the optic fiber of the optrode.

For each electrode location, 5 ms and 50 ms single light pulses were delivered at 2 mW. Following offline sorting of waveforms, ChR2-expressing CRH single units were identified by their time-locked responses to light (see details in *Data analyses and statistics: Identification of light-evoked spikes*). Specifically, we clustered all waveforms (both spontaneous and light-evoked in a blinded manner) for single unit isolation (L-ratios < 0.05). Thereafter, single units clustered with the light-evoked waveforms were considered as light responsive and thus CRH neurons, and other single units simultaneously recorded with light responsive neurons but did not respond to the light were considered as light non-responsive, which includes both non-CRH neurons and CRH neurons with insufficient ChR2 expression and/or light exposure.

### Sciatic nerve stimulation

Under isoflurane anaesthesia, a small incision was made into the hind limb of the animal and the sciatic nerve was isolated from surrounding tissue. A bipolar tungsten stimulating electrode was gently placed on the nerve and the nerve was kept hydrated with periodic applications of sterile saline. The contralateral nerve to the recording site was stimulated with 1.6 mA negative pulses at 20 Hz (5 x 0.5 ms) using a pulse stimulation unit (S88, GRASS Instrument Co) connected to a stimulus isolation unit (Model PSIU6, GRASS Instrument Co) triggered via TTL signals.

### Histology

At the end of recording, animals were euthanized with an overdose of pentobarbital sodium (150 mg/kg, i.p.) and transcardially perfused with 0.9 % saline and 4% paraformaldehyde (PFA). The brain was collected and left to post-fix in 4% PFA at 4 °C overnight. The brain was washed in phosphate buffer saline (PBS) solution and sliced into 40 µm coronal sections and counter stained with 4,6-diamidino-2-phenylindole (DAPI). The slices were mounted and verified for CRH expression (tdTomato), ChR2 expression (eYFP), and the dye-painted electrode tract (Vybrant DiD, Thermo Fisher V22889).

### Experimental design

Each recording started with the baseline recordings of spontaneous firing without any intentional sensory stimulation for at least 10 minutes. This was followed by light stimulation for CRH neuron identification, then subsequently sciatic nerve stimulation (minimum 3 minutes between light stimulation and sciatic nerve stimulation).

### Data analyses and statistics

Data analysis were carried out using built-in and custom-built software in MATLAB (MathWorks). Graphs were made and statistical analysis were conducted using Prism 8 (GraphPad).

#### Identification of light-evoked spikes

Time-locked response to light stimulation was tested by an increase in spiking activity to a 5 ms pulse of blue light (25 trials). The response was plotted on a peristimulus time histogram (PSTH) aligned to the onset of light. Peristimulus frequency before and after light onset (20 ms windows) was compared by a trial-by-trial paired t-test: a significant increase was defined as light-responsive. The event probability before and after light (20 ms windows) was calculated as the number of trials that had at least 1 event within the response window, divided by the total number of trials.

#### Spike pattern analysis

For each single-unit, spike activity was classified into bursts and single spikes. Bursts were detected as series of two or more spikes starting with the initial inter-spike interval (ISI) less than 6 ms and subsequent ISIs less than 20 ms. Although rare, to prevent contamination of bursts by high-firing single spikes which gradually speed up (*i.e.* shortening ISIs) into the burst range, we adopted a secondary criterion for bursts to be preceded by an ISI >25 ms. Spikes not associated with bursts were defined as single spikes. For analyses involving burst rates (*i.e.* burst rate, burst index, inter-burst interval (IBI)), each burst was counted as a single event regardless of burst length (the number of spikes per burst). IBI distribution was first calculated for each cell (100 ms bins, plotted by upper limit of bin), and then the group distribution was generated by plotting the mean and standard deviation. Similarly, the distribution of burst length (the number of spikes per burst) was first calculated for each cell, and then the group distribution was generated by plotting the mean and standard deviation.

To examine spiking dynamics that may influence burst activity, the probability of burst initiation (ISI_(*i*)_ <6 ms) as a function of preceding silence (ISI_(*i-1*)_) was plotted for each cell. To do this, spikes were binned by ISI_(*i-1*)_ (10 ms and 500 ms bins for periods <500 ms and ≥500 ms, respectively), and the probability of ISI_(*i*)_ <6 ms was calculated for each bin for each cell. Then, the group average was plotted with the mean and standard deviation. To examine the influence of preceding silence on burst length, spikes were binned by event length, and the mean preceding silence was calculated for each cell. Then, the group average was plotted with the mean and standard deviation. The silent period analysis was repeated for proceeding silent periods. In this case, the probability of burst (ISI_(*i*)_ <6 ms) as a function of the proceeding silence (ISI_(*i+1*)_) was plotted. Finally, the mean proceeding silence was compared by event lengths.

#### Firing rate analysis

The average baseline firing rates were calculated as the total number of spikes divided by total time during the spontaneous baseline recording (10 min). The firing rate time course was calculated as the number of spikes in every 10-second bins. The running average firing rate represents a moving average of five consecutive bins (bin(*i-2*) + bin(*i-1*) + bin(i) + bin(*i+1*) + bin(*i+2*) / 5) and was used to visualize spontaneous fluctuations of firing rate. These fluctuations in firing rate were then overlaid onto ISIs plotted across time to visualize the relationship between firing rate to firing patterns. To examine the relationship between burst rate and total spike rate, for every 10-second bin in individual units, total spike rate was plotted against burst rates of 0.1, 0.2, 0.3, 0.4, 0.5 or >=0.6 bursts/s. The relationship was plotted for individual unit (**Fig. 3L**), as well as pooled across units (**Fig. 3M**, n = 18).

#### Sciatic nerve stimulation

The response to sciatic nerve stimulation was plotted on PSTHs aligned to the onset of stimulation (the first of the 5-pulse train). A paired t-test was performed between the average post-onset firing rates (2 seconds across 30-60 trials) and the baseline firing for each cell. Responses to sciatic nerve stimulation was analysed in three categories: all spikes regardless of spiking pattern, single spikes only, and bursts only (each burst train counted as a single event regardless of the burst length).

### Recurrent inhibitory network model

*Single neuron model fit.* Single neurons were modelled using the Adaptive Exponential Integrate and Fire (AdEx) model (Brette and Gerstner, 2005).

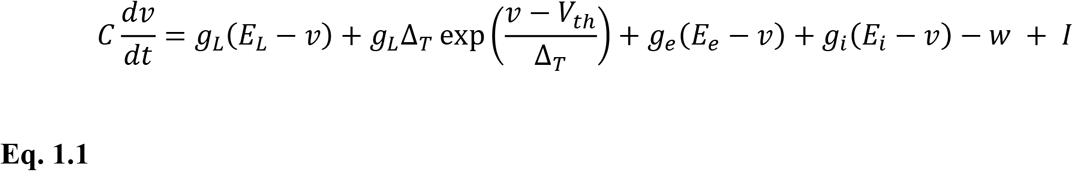

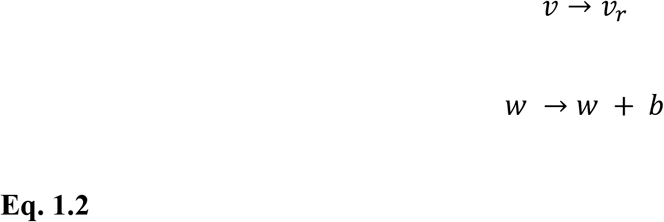

Here, *C* is the membrane capacitance, *w* is an adaptation variable, *I* is the applied current, *g_L_* is the leak conductance, *E_L_* is the resting membrane potential, Δ*_T_* is the slope factor, and *V_th_* is the threshold potential. The slope factor determines the sharpness of the threshold. The membrane potential *v* is modelled as a sum of these conductance parameters (**Eq. 1.1**). When the membrane potential *v* exceeds the threshold *v_th_* the neuron emits a spike, and the neuron is reset according to **Eq. 1.2**. *w* is computed as a function of the subthreshold, and spike-triggered adaptations, respectively (**Eq 1.3**)

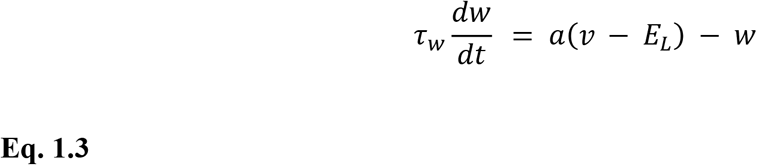

where *τ_w_* is the time constant and *a* is subthreshold adaptation conductance.

To refine the model to capture the behaviour of CRH_PVN_ neurons, we fit the model with whole-cell patch-clamp recordings of CRH_PVN_ neurons in acute brain slices (n = 33, 5 mice) using methods detailed in (Matovic et al., 2020). Recordings consisted of 14 trials containing a hyperpolarizing (-20 pA for 200 ms) pulse followed by a stepped depolarizing current pulse (10 pA steps for 500 ms). The same stimulus was used in subsequent model simulations. First, the firing rate-current (F-I) curve for the model was simulated across a large parameter space. Then, the posterior distribution for the parameter space was fit using the Sequential Neural Posterior Estimation (SNPE) method as previously described (Gonçalves et al., 2020). Second, for each reference cell, the F-I curve was used to draw a restricted parameter space from the previously fit posterior. Then, 200 rounds of evolutionary optimization were used to refine the model (Lynch and Houghton, 2015). The penalty (or evaluation of fit) was calculated as the sum of the mean squared error (MSE) between the subthreshold voltage of the model and reference, and the log MSE between the F-I curve of the model and reference. Models that produced an MSE above 5 mV (for the subthreshold voltage) and above 10 Hz (for the F-I curve) were considered poor fits, and were discarded. To produce a robust general model, the average of the parameter fit for all reference traces was used in the network model (TABLE 1)

#### Network model

To construct the network model, we utilized a simplified recurrent network template with two distinct populations of AdEx neurons (Destexhe, 2009). These neurons consisted of a modified AdEx model with added excitatory and inhibitory synaptic parameters **(Eq. 1.1**).

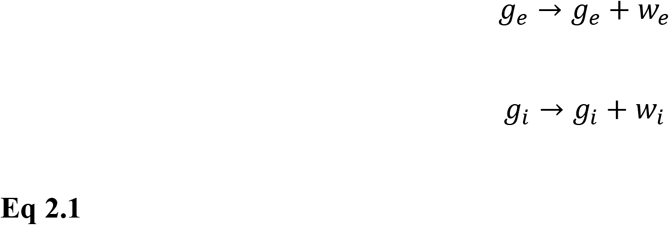

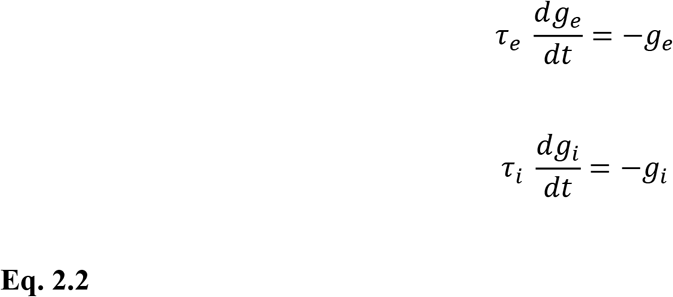

Here, *g_e_* & *τ_e_* and *g_i_* & *τ_i_* represent the inhibitory and excitatory conductance and time constant, respectively. Following a presynaptic spike, the inhibitory and excitatory conductance were incremented according to **Eq. 2.1.** Followed by a loss of conductance via a simple exponential decay (**Eq. 2.2**).

Both populations were interconnected, with one population presenting as excitatory and one as inhibitory. For simplicity, only the excitatory neurons received external input. First, we constrained the excitatory portion (N=500) of the network to our biologically realistic CRH_PVN_ neurons. To do so, we used previously fit parameters from the single neuron model as described in *Single neuron model fit*. Next, the remaining inhibitory population was constrained with parameters, reflecting the tonic spiking GABAergic neurons of the thalamus (Destexhe, 2009). Changes in adaptation current *b* were modelled with instantaneous increment followed by an exponential decay function (**Eq 3.1**), reflecting the slow change in intracellular parameters.

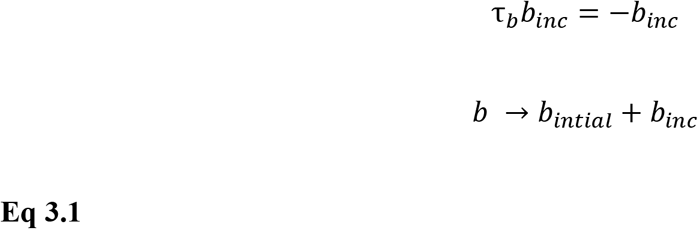

For each synapse, the probability of release *p* was modelled by the logistic differential equation

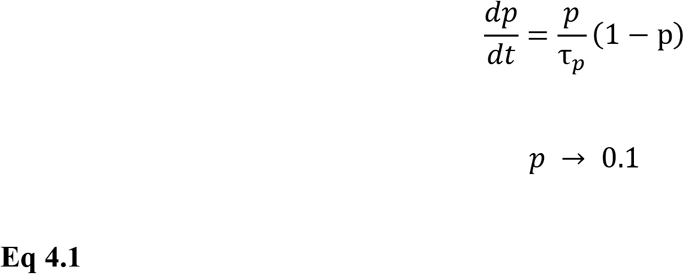

Based on previous literature, the excitatory input of CRH_PVN_ neurons to the GABAergic population was modeled to be mediated by the CRHR1 receptor (Jiang et al., 2018, 2019; Ramot et al., 2017). Therefore, this synapse was modelled by a slow-growing alpha function, a common model of neuromodulator function (Destexhe et al., 1994):

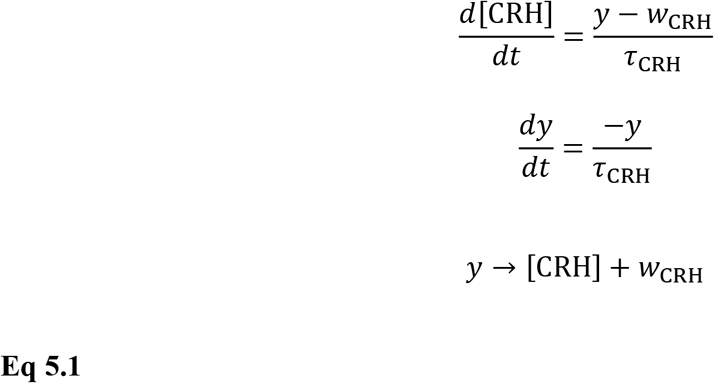

The remaining free parameters *w_e_*, *w_i_*, *τ_e_*, *τ_i_*, *τ*_CRH_, and the EPSP input frequency were tuned with respect to the observed extracellular spike trains *in vivo*. To begin with, we selected a representative opto-tagged CRH neuron baseline recording as reference (**Fig. 2B**). Next, we selected two characteristic (50 seconds) sections of the reference spike train that reflected bursting and single spiking independently. Then, several rounds of evolutionary algorithms were used to tune the parameters. For each round, the network was simulated with the given parameters with an increased adaptation current (2 x 50 s). Then, the distance between simulated and reference data was computed as the earth mover’s distance (Wasserstein metric) between the log-scaled ISI distributions (Olkin and Pukelsheim, 1982). The total distance was taken as the sum of the smallest 100 distances between each simulated unit and the reference. The standard adaptation current and the increased adaptation current simulations were compared to the bursting and tonic references, respectively.

#### Generation of network clamp input for ex vivo experiments

To investigate the ability of our model to induce burst in biological neurons *ex vivo*, we extracted the input currents from a model neuron in the network. First, a model neuron, isolated within the network, had its initial resting membrane potential V_mØ_ and leak reversal E_L_ constrained to the mean resting membrane potential of the CRH_PVN_ neurons in our recording conditions *ex vivo* (-67.9mV, **Table 1**). Then, the model was simulated for 50 seconds, and the model neuron was allowed to respond to the incoming input freely. Every time-step (100 µs), the sum of the incoming excitatory, inhibitory, and adaptation currents were extracted following **Eq. 6.1**:

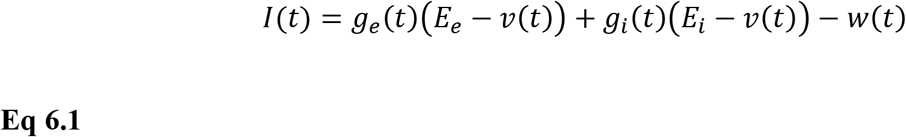

Where *g_e_, g_i_*, *v*, and *w* are extracted from the values of the individual neuron found in **Eq 1.1** at given time step *t*. In essence, this formulation allowed observation of the model neuron state without intervention. We theorized that inclusion of the computed leak current: *g_L_* (*E_L_* − *v*), was unnecessary as the *ex vivo* neurons express this intrinsically. Our network model generated very large synaptic currents (∼500 pA), which is unlikely to reflect realistic inputs to CRH_PVN_ neurons observed in *ex vivo* preparation (Salter et al., 2018; Sterley et al., 2018; Wamsteeker Cusulin et al., 2013). Thus, we capped the maximum amplitude of PSCs to ±200pA without changing the timings of PSPs.

#### Software

Simulations and modelling were completed using software packages within the Python 3.5 ecosystem. Single neuron and network simulations were completed using the Brian 2 (2.4) package (Stimberg et al., 2019). MSE, EMD, and other mathematical operations were computed using modified implementations of the functions available in NumPy (1.20.1) and SciPy (1.6.0) software packages (Harris et al., 2020; Virtanen et al., 2020). Backend probability inference was completed using the sbi (0.16.0) package (Gonçalves et al., 2020). Optimization of fits was performed using the nevergrad (0.4.3) package (Liu et al., 2020). Specifically, we utilized a custom portfolio consisting of two-points differential evolution (two-points DE), covariance matrix adaptation evolutionary strategy (CMA-ES), and a random search using scrambled Hammersley sequence (SCR-Hammersly). Axon binary file (ABF) recordings were imported to python using the pyABF package (Harden, 2020). Extraction of single-neuron action potentials and other electrophysiological features from intracellular recordings was performed using the ipfx (0.1.1) package (Gouwens et al., 2019). A generalized formulation of the fitting code, including the interface between Brian 2, nevergrad, and sbi will be made public on GitHub once the paper is accepted. This includes the methodology for comparing EMD between spike trains in 1 and 2 dimensions using a sliced Wasserstein implementation.

## ACKNOWLEDGEMENT

We thank Dr. Karl Deisseroth (Stanford University) for providing pAAV-EF1a-double floxed-hChR2(H134R)-EYFP-WPRE-HGHpA was a gift from (Addgene plasmid # 20298 ; http://n2t.net/addgene:20298 ; RRID:Addgene_20298). This work was funded through a Discovery Grant (RGPIN-2015-06106) from the Natural Sciences and Engineering Research Council of Canada (NSERC; to W. I.), a Project Grant (PJT-148707) from the Canadian Institutes of Health Research (CIHR; to W. I.), BrainsCAN at Western University through the Canada First Research Excellence Fund (CFREF; to W. I and L. M), and the Compute Canada (to L. M.). A. I was a recipient of an Ontario Graduate Scholarship, S. M. is a recipient of the Canadian Open Neuroscience Platform studentship.

## COMPETING INTERESTS

Authors have no competing interests to declare.

---

**Figure 2-Figure Supplement 1.**
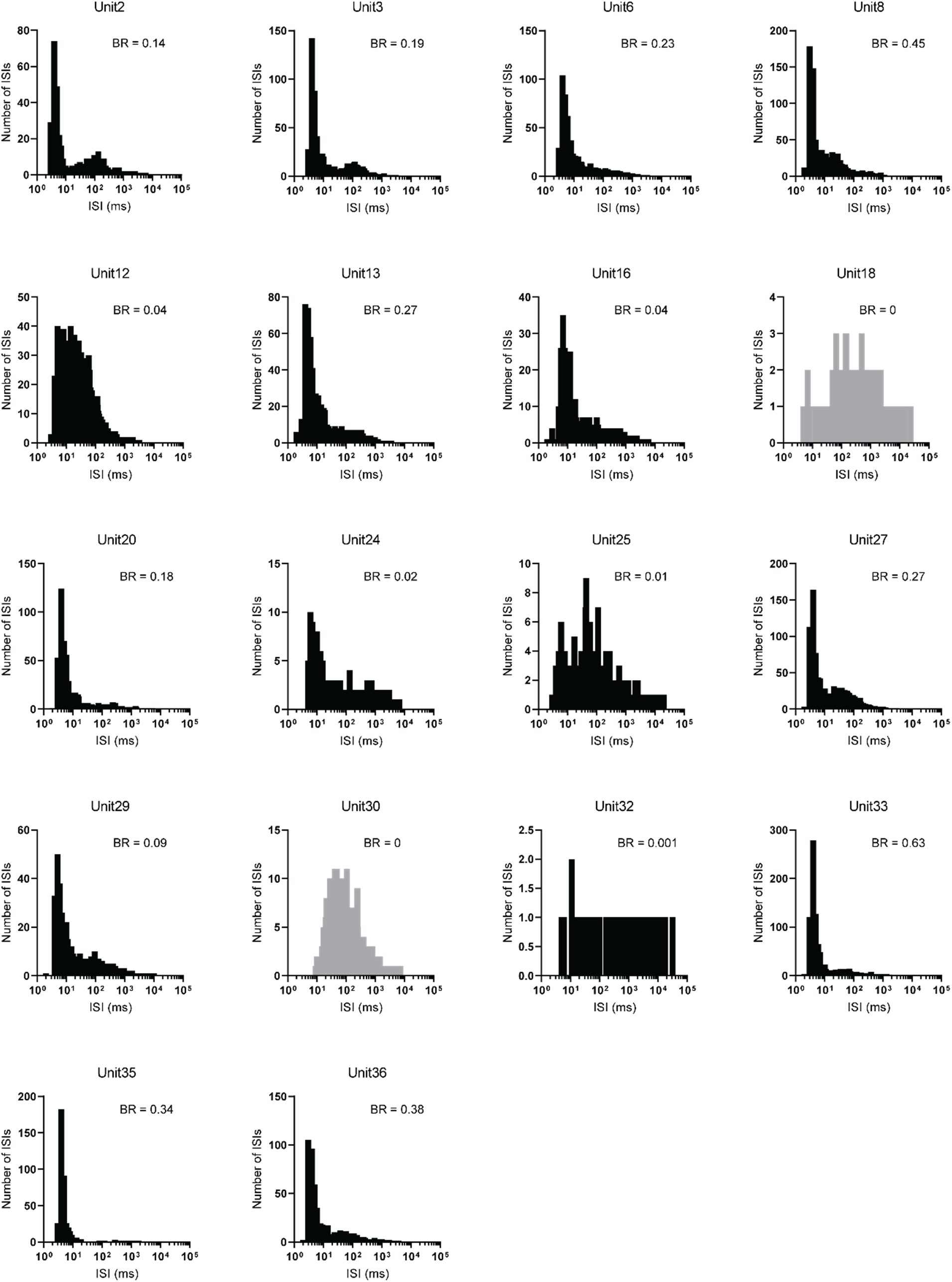
Inter-spike interval (ISI) histograms for all CRH_PVN_ units. Burst rate (BR) is indicated on the top-right corner of individual histograms. Unit 18 and 30 did not fire bursts and are shown in gray.

## Notes

### Competing Interest Statement

The authors have declared no competing interest.

